# Amnion signals are essential for mesoderm formation in primates

**DOI:** 10.1101/2020.05.28.118703

**Authors:** Ran Yang, Alexander Goedel, Yu Kang, Chenyang Si, Chu Chu, Yi Zheng, Zhenzhen Chen, Peter J. Gruber, Yao Xiao, Chikai Zhou, Nevin Witman, Chuen-Yan Leung, Yongchang Chen, Jianping Fu, Weizhi Ji, Fredrik Lanner, Yuyu Niu, Kenneth Chien

## Abstract

Essential genes for murine embryonic development can demonstrate a disparate phenotype in human cohorts. By generating a transcriptional atlas containing >30,000 cells from postimplantation non-human primate embryos, we discovered that *ISL1*, a gene with a well-established role in cardiogenesis, controls a gene regulatory network in primate amnion. CRISPR/Cas9-targeting of *ISL1* resulted in non-human primate embryos which did not yield viable offspring, demonstrating that *ISL1* is critically required in primate embryogenesis. On a cellular level, mutant *ISL1* embryos displayed a failure in mesoderm formation due to reduced BMP4 signaling from the amnion. Via loss of function and rescue studies in human embryonic stem cells we confirmed a similar role of *ISL1* in human *in vitro* derived amnion. This study highlights the importance of the amnion as a signaling center during primate mesoderm formation and demonstrates the potential of *in vitro* primate model systems to dissect the genetics of early human embryonic development.

## Introduction

Studies in genetically modified model organisms, in particular the mouse, have allowed us to study mammalian embryonic development in astonishing detail and have laid the foundations to understanding human development. However, in certain cases, the phenotype observed in knock-out mouse models differs from observations in human cohorts. The LIM-domain transcription factor *ISL1* has a well-established role in mammalian cardiac development and is expressed in multipotent cardiovascular progenitor cells in mice ^1–3^ and humans ^4, 5^. In line with this, *Isl1* loss of function mice have severe cardiac defects leading to embryonic lethality at E10.5 ^6, 7^. Despite its established role in heart development, loss of function variants in the *ISL1* locus have rarely been associated with cardiac defects in humans and are underrepresented in large human cohorts of congenital heart malformations like the Pediatric Cardiac Genomics Consortium (PCGC) ^8, 9^. In detail, among the 23,000 alleles reported in the PCGC cohort, 112 *ISL1* variants have been identified, none of which were damaging *de novo* mutations. Based on this low frequency of damaging *ISL1* variants we hypothesize that *ISL1* has an alternative, essential requirement during early primate embryogenesis.

Studies of *in vitro* cultured human embryos have shown, that *ISL1* is not expressed in the preimplantation blastocyst ^10^. One of the key steps during mammalian development following implantation is the formation of the three primary germ layers. This occurs in a complex process termed gastrulation where cells from the columnar shaped epiblast undergo epithelial-to-mesenchymal transition and move ventrally and anteriorly to form the mesodermal cells ^11–13^. It is believed that improper gastrulation occurs frequently in human embryos and accounts for a significant proportion of early miscarriages in the human population.

The tight regulatory network governing this process has been well studied during murine embryonic development ^11, 14^, but is largely elusive in humans. Recently, two publications on cynomolgus embryogenesis ^15, 16^ and one publication on human embryogenesis ^17^ have created a framework of this developmental time window in primates and characterized the major cell populations involved in gastrulation. However, their interplay and the transcriptional networks guiding this essential step remain unknown.

Here, we created a high-resolution map of the peri-gastrulation development of NHP embryogenesis. We identify an *ISL1*-dependent gene regulatory network that is specifically active in amnion. Disturbance of this network in NHP embryos by CRIPSR/Cas9-mediated gene-editing of *ISL1* led to embryonic lethality due to significant downregulation of BMP4 signaling from the amnion and subsequent failure to form mesoderm. We confirmed these findings in a microfluidic-based embryonic sac model of amnion-epiblast interactions using *ISL1*-null human embryonic stem cells suggesting that these findings also apply to humans. Taken together, this study demonstrates a novel, primate-specific role of *ISL1* in early embryogenesis and shows for the first time that signals from the amnion are indispensable for mesoderm formation in primate embryos.

## Results

### Loss of *ISL1* leads to embryonic lethality

To assess whether *ISL1* plays a functional role in primate embryogenesis, we generated *ISL1* hypomorphic mutant NHP embryos through one-cell stage CRISPR/Cas9 injections with gRNAs creating a long deletion in the *ISL1* locus and transferred them into NHP surrogate mothers (Fig. 1a). Genotyping of the mutant embryos revealed a mosaic pattern with presence of ind/dels of different sizes in the targeted region of the *ISL1* locus and no alterations in selected off-targets from the in-silico prediction ^18^ (Supplementary Fig. 1). The frequency of mutations resulting in an internally shortened, but functional ISL1 was significantly higher in the transcriptome (Supplementary Fig. 1a), most likely due to nonsense-mediated decay of mutated *ISL1* message.

**Fig. 1:**
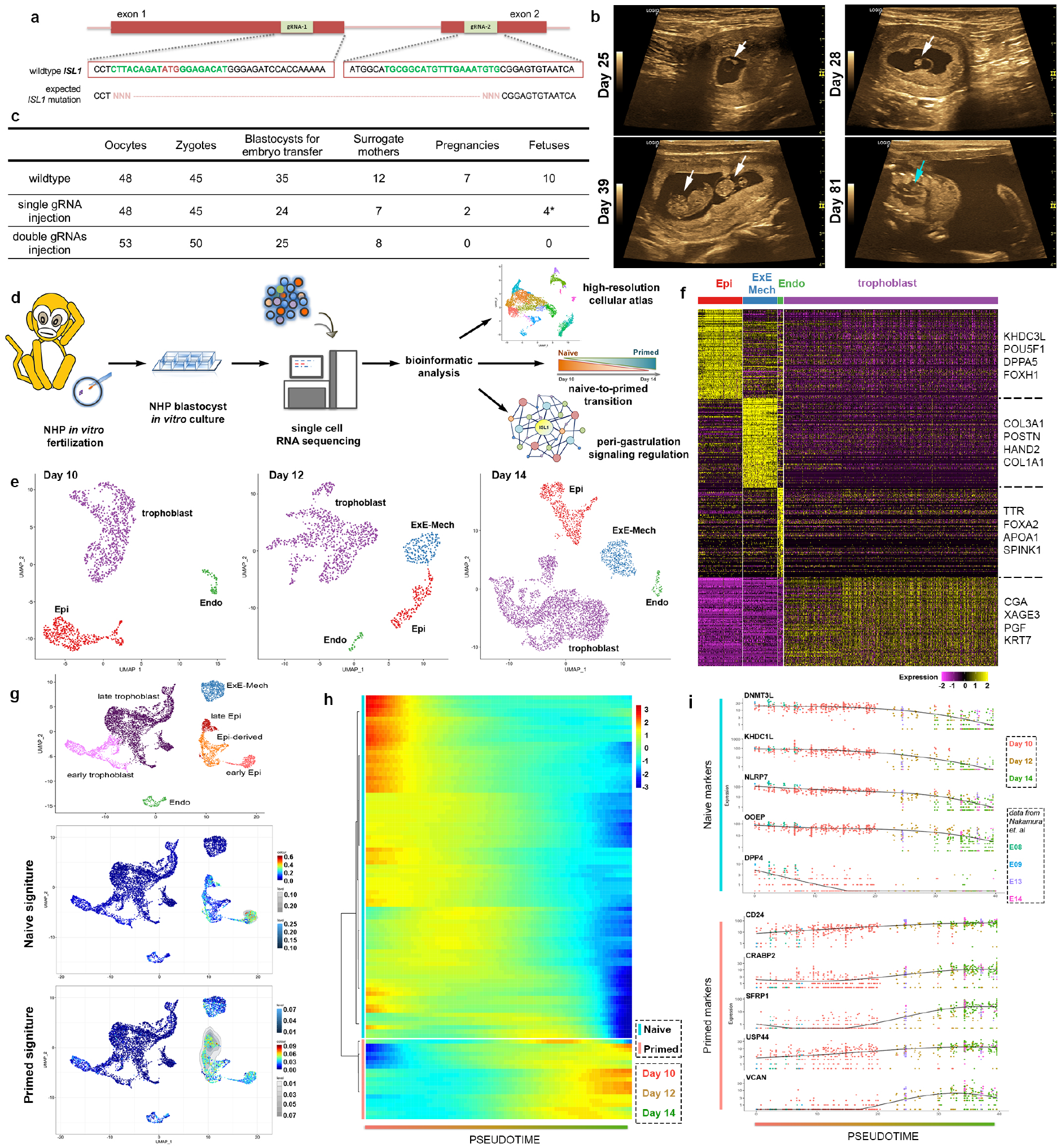
High resolution transcriptomic map of peri-gastrulation events in wildtype *in vitro* cultured NHP embryos. See also Supplementary Fig. 1-3. **a,** guide RNA design for generating ISL1 mutation. **b**, ultrasound scanning for pregnancy diagnosis. Days correspond to time after fertilization. White arrows indicate fetus, and blue arrow fetal heart. **c**, summary of embryo transfer. Asterisk indicates wildtype fetuses. **d**, scheme of the workflow. **e**, UMAP plot of all cells from the *in vitro* cultured embryos at the different time points (Day 10, Day 12 and Day 14) colored by cell type. **f**, heatmap showing the scaled expression at Day 14 of 100 differentially expressed genes (DEGs) for each cell type identified in panel selected by adjusted p-value. **g**, UMAP plot of all cells of the integrated dataset from the *in vitro* cultured embryos (Day 10, Day 12 and Day 14) with cells from the *in vivo* NHP embryos (E08, E09, E13 and E14) colored by cell type. Visualization of the single cell signature score reflecting the naïve and the primed state of pluripotency on the UMAP plot of the integrated dataset. **h**, heatmap showing the scaled expression of 100 DEGs across pseudotime in epiblast development selected by the Moran’s I statistics. Unsupervised clustering of genes into a blue cluster containing genes downregulated over pseudotime and a red cluster containing genes upregulated over pseudotime. **i**, gene expression of 5 exemplary genes downregulated over pseudotime (blue cluster) and 5 exemplary genes upregulated over pseudotime (red cluster) from (**g**). Black line showing the fitted expression trend for each gene over pseudotime. Epi, epiblast; ExE-Mech, extraembryonic mesenchyme; Endo, endoderm.

Pregnancy of NHP surrogate mothers was assessed by the presence of an embryonic structure within a gestation sac on ultrasound imaging from 4 weeks of gestation (Fig. 1b). The pregnancy rate per NHP surrogate mother after transfer of mutant embryos was 0% as compared to 58.3% with wildtype embryos (Fig. 1c). Transfer of embryos that were targeted with injection of only a single gRNA, leading to a slightly lower mutation rate, resulted in a pregnancy rate of 28.6% (Fig. 1c). Strikingly, genotyping of all 4 fetuses from this experiment showed an unmodified *ISL1* locus on both alleles, confirming the suspected early requirement of *ISL1* for proper embryonic development.

### Single cell map of post-implantation NHP embryos

To map the expression of *ISL1* in the early embryo we created a high-resolution transcriptomic atlas by single cell RNA (scRNA) sequencing of 11 *in vitro* cultured cynomolgus macaque embryos at three different time points (Day 10, Day 12 and Day 14) (Fig. 1d-e). 7194 cells passing quality control (Supplementary Fig. 2) were embedded for each day separately in lowdimensional space (Fig. 1e and Supplementary Fig. 3a). In line with previous results ^15, 16^, the cells grouped into four main cell types, namely trophoblast, endoderm, epiblast with its derivatives and extraembryonic mesenchyme (Fig. 1e-f). Integration of our dataset with a published scRNA sequencing dataset of *in vivo* cynomolgus embryos ^19^ (Fig. 1g, Supplementary Fig. 3b) revealed a striking difference in the transcriptomic profile between early (Day 10 + E08/E09) and late (Day 12/Day 14 + E13/E14) epiblast, reflecting the transition from a naïve to a primed state, which has been suggested before to happen during this time window ^20^. Indeed, a published gene signature of naïve human embryonic stem cells (hESCs) ^21^, was highly enriched in the early peri-implantation epiblast at Day 10 (Fig. 1g), while genes belonging to the primed hESCs signature were enriched in the late epiblast at Day 12 and 14 (Fig. 1g). Aligning cells from the early and late peri-implantation epiblast in pseudotime disclosed a set of differentially regulated genes that formed two distinct clusters based on their expression dynamics (Fig. 1h). Genes previously associated with a naïve state ^20–22^, including *DNMT3L, KHDC1L, NLRP7, OOEP* and *DPP4* were significantly downregulated over pseudotime (Fig. 1i). In contrast, genes associated with a primed state ^22–24^, including *CD24, CRABP2, SFRP1, USP44* and *VCAN* showed strong upregulation (Fig. 1i). This expression pattern was observed in cells from our dataset as well as in cells from the *in vivo* dataset, suggesting that the naïve to primed transition happening *in vivo* can faithfully be recapitulated in *in vitro* cultured embryos.

### Gene regulatory network of early epiblast

Taking advantage of our high-resolution scRNA map, we next investigated the appearance of epiblast-derived cell populations. We identified cells expressing *ISL1* alongside with amnion marker genes such as *TFAP2A* ^15^ as early as Day 10 (Fig. 2a and Supplementary Fig. 3c-d). Embryos at this stage also consisted of a relatively large cell population expressing genes typical of a mesodermal signature such as *MIXL1* and *MESP1* (Supplementary Fig. 3e), which were previously annotated as early gastrulating cells ^15^. However, in addition to their mesodermal signature, these cells show high expression of the transcription factor *ETS1* and the cell adhesion protein *PODXL* (Supplementary Fig. 3f), which mark extraembryonic mesoderm in mice ^25^, while showing low levels of expression of the receptor tyrosin-kinase *EPHA4* or the transcription factor *ZIC3* (Supplementary Fig. 3g) expressed preferentially by murine embryonic mesoderm ^25^. This suggests, that the epiblast-derived mesodermal-like cells present in Day 10 embryos are likely extraembryonic mesoderm, which seem to appear prior to gastrulation. Extraembryonic mesenchyme, a cell population unique to primates that contributes to a number of extraembryonic tissues ^19, 26–28^, was first present in Day 12 embryos (Fig. 2a). The close proximity of extraembryonic mesoderm and mesenchyme in the UMAP plot (Fig. 2a) as well as shared expression of genes such as *PODXL* and *ETS1* (Supplementary Fig. 3h), advocate that extraembryonic mesoderm develops into extraembryonic mesenchyme. The large increase in cell number in the extraembryonic mesenchyme over a few days could indicate that this cell population gets additional contributions from other parts of the embryo, such as the trophoblast or the endoderm, as previously suggested ^29^, although we did not find any evidence for this on the transcriptional level.

**Fig. 2:**
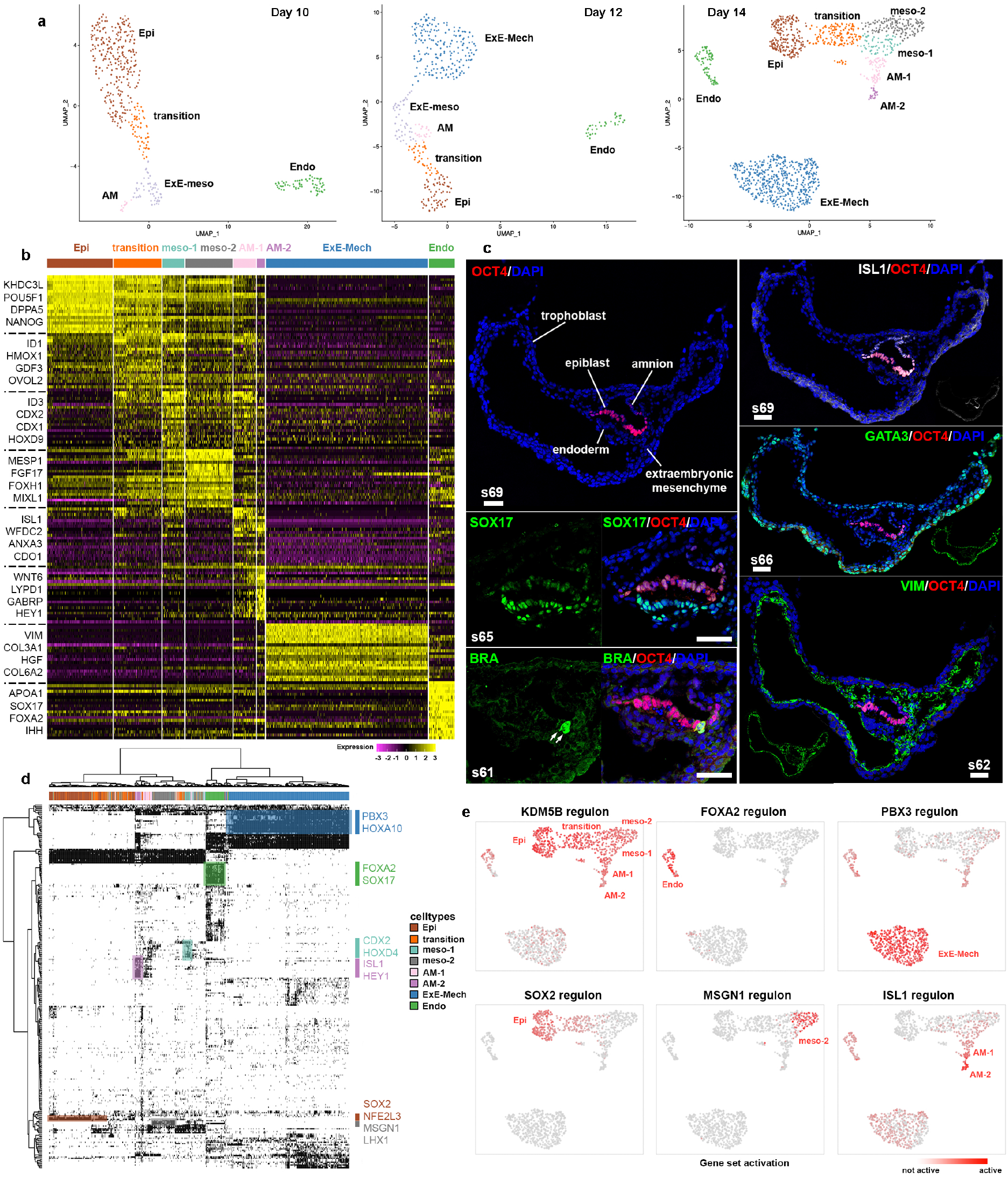
Gene regulatory networks in epiblast and its derivatives, endoderm and extraembryonic mesenchyme. See also Supplementary Fig. 3. **a**, UMAP plot of cells from the in vitro cultured embryos at the different time points (Day 10, Day 12 and Day 14) mapping to the Epi and its derivatives, Endo and ExE-Mech colored by cell type. **b**, heatmap showing the scaled expression at Day 14 of 20 DEGs for each cell type identified in (**a**) selected by adjusted p-value. **c**, fluorescent staining of major cell types found from scRNA sequencing. Scale bar 50 μm. White number indicates section number. **d**, binary activity matrix of regulons identified at Day 14 by gene regulatory network interference active in at least 1% of the cells clustered unsupervised. Selected master regulators are depicted in the color corresponding to the cell type they show activity in. **e**, gene set activity of the selected regulons at Day 14 in the different cell types depicted on the UMAP plot from (**a**). Epi, epiblast; Endo, endoderm; ExE-Meso, extraembryonic mesoderm; ExE-Mech, extraembryonic mesenchyme; AM-1, amnion 1; AM-2, amnion 2; meso-1, mesoderm 1; meso-2, mesoderm 2.

In Day 14 embryos we identified two clusters of mesodermal cells (Fig. 2a-b). Cells in cluster mesoderm 1 preferentially expressed members of the caudal gene family *CDX1, CDX2* and *CDX4*, while cells in the cluster mesoderm 2 were expressing relatively higher levels of *MESP1* and *GSC* (Fig. 2b, Supplementary Fig. 3i-j). Cells in the cluster Amnion 2 showed high expression of amnion markers such as *WNT6, GABRP*, and *ISL1* (Supplementary Fig. 3k). Transcripts of these markers, although at lower levels, were also detected in the cluster Amnion 1 alongside with mesoderm-associated genes such as *CDX1* and *CDX2* (Fig. 2b, Supplementary Fig. 2i-k). The amnion markers *WNT6, GABRP* and *ISL1* were enriched in amniotic cells across all cell types of the embryo while other markers such as *HEY1* were also expressed in subpopulations of the trophoblast (Supplementary Fig. 3l).

Immunofluorescent imaging of sectioned *in vitro* cultured embryos at Day 14 confirmed the presence of the different cell populations identified in the scRNA sequencing analysis (Fig. 2c). In particular, we observed specific staining for ISL1 in amniotic cells (upper right panel) overlying the epiblast staining positive for OCT4 *(POU5F1*, upper left panel). Endoderm showed strong signal for SOX17 (mid left panel), while trophoblast stained positive for GATA3 (mid right panel). The presence of Brachyury (BRA, *TBXT)* positive cells located ventral of the embryonic disc (lower left panel) suggests that the mesodermal cells identified in the scRNA sequencing data at Day 14 are indeed emerging mesoderm during gastrulation. Extraembryonic mesenchyme, staining positive for Vimentin *(VIM)* was clustering around the embryonic disc, in particular in the region which will most likely develop into the connecting stalk in later stages, but was also lining the entire trophoblast (lower right panel).

Cell type specification is under the control of transcription factors that bind to cis-regulatory regions, forming gene regulatory networks (GRN) ^30^. GRN analysis at Day 14 using SCENIC (Single-cell regulatory network inference and clustering) ^31^ revealed sets of GRNs specifically active in these populations (Fig. 2d-e). The histone demethylase *KDM5B*, creating bivalent histone marks during development ^9^, was identified in this analysis to control a GRN active in epiblast and all its derivatives, while the pluripotency factor *SOX2* ^24^ controlled a network specifically active in the epiblast (Fig. 2e). One of the most active GRNs in amniotic cells was an ISL1-dependent network (Fig. 2e), signifying that *ISL1* is not only a specific marker of the amniotic cell population in primates, but could be functionally important. This finding is in sharp contrast to the mouse, where *Isl1* is first expressed in cardiac progenitor cells of the lateral plate mesoderm, but absent from the early embryo before E7.0 ^7, 32^.

### *ISL1* hypomorphic mutant embryos fail to form mesoderm

To investigate the functional role of this ISL1-dependent GRN in post-implantation development, the *ISL1* hypomorphic mutant embryos were cultured *in vitro.* Mutant embryos developed normally up to Day 10 (Fig. 3a and Supplementary Fig. 4a). After Day 12 they progressively lost structure and at Day 14, the embryonic disk was no longer distinguishable (Fig. 3a and Supplementary Fig. 4a). This was reflected in the integrated analysis of 26136 cells from Day 10, Day 12, and Day 14 mutant embryos (Supplementary Fig. 2) with the wildtype dataset by a progressive loss of cells in the Epi-derived cluster while the other lineages were preserved (Fig. 3b and Supplementary Fig. 4b-c). Subclustering of the Epi-derived cells at Day 14 showed a drastic reduction of cells in the mesodermal clusters in the mutant embryos accompanied by an overrepresentation of amniotic cells (Fig. 3c). In line with this, the number of cells expressing the mesoderm markers *TBXT, EOMES, MIXL1, MESP1*, and *CDX2* ^33–35^ was reduced in the mutant embryos among epiblast derivatives (Fig. 3d) as well as across the whole dataset (Supplementary Fig. 4d).

**Fig. 3:**
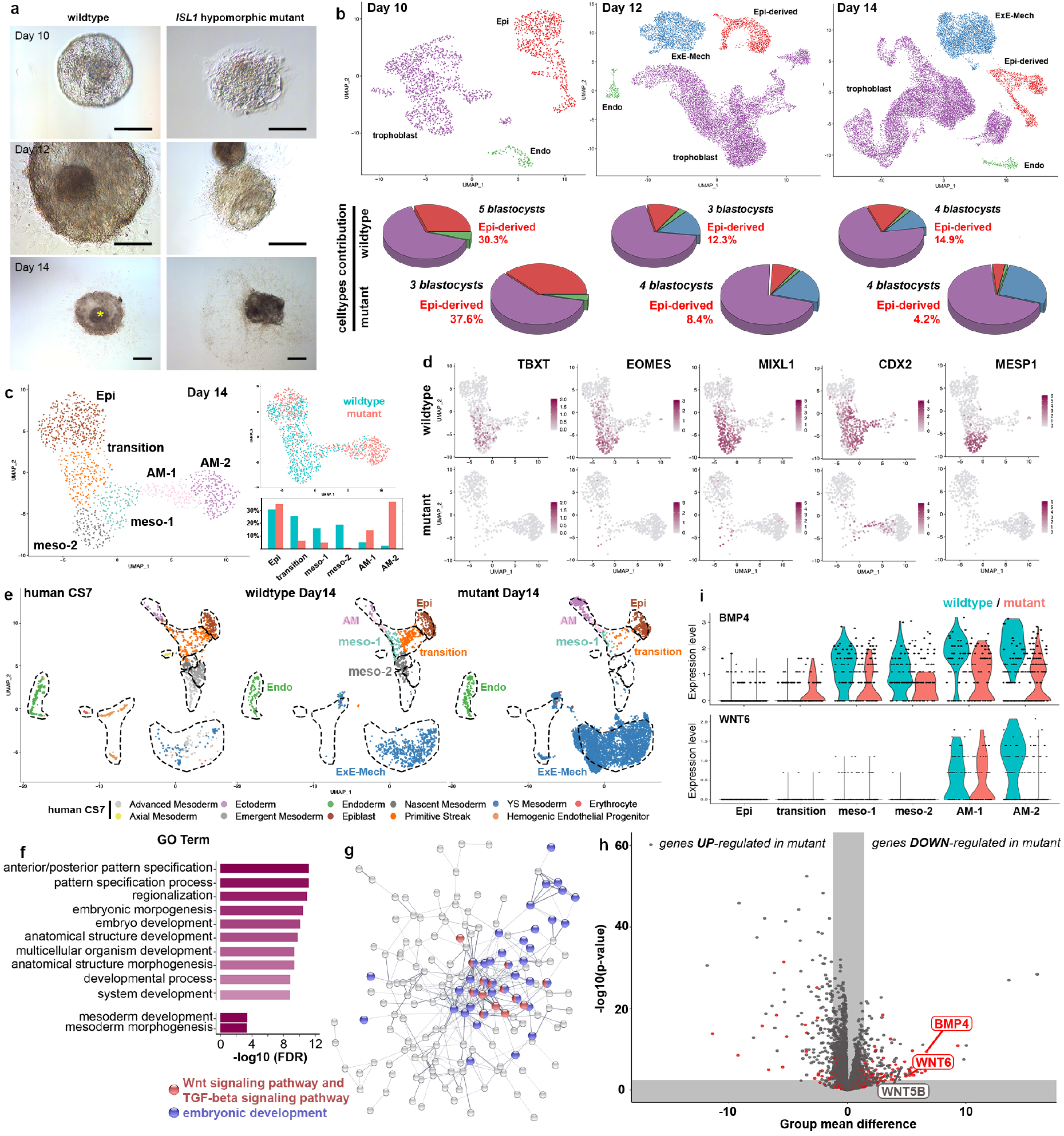
*ISL1* hypomorphic mutants fail to form mesoderm. See also Supplementary Fig. 4. **a**, the morphology of wildtype (left) and *ISL1* hypomorphic mutant (right) NHP embryos at Day 10, Day 12 and Day 14. Scale bar, 200 μm. Asterisk indicates the embryonic disk. **b**, UMAP plot of cells mapping to the epiblast and its derivatives from the integrated dataset of wildtype and mutant embryos at the different time points (Day 10, Day 12 and Day 14) colored by cell type. Pie charts indicate the relative contribution of cells. **c**, UMAP plot of cells mapping to the epiblast and its derivatives at Day 14 and the relative contribution of cells to the various cell types in wildtype (blue) and mutant (red) embryos. **d**, Expression of mesodermal marker genes in cells from the epiblast and its derivatives of Day 14 wildtype (top) and mutant (bottom) embryos. **e**, UMAP plot of the integrated dataset of all cells despite trophoblast from the *in vitro* cultured embryos (Day 10, Day 12 and Day 14) with cells from the human Carnegie stage 7 gastrula colored by cell types. **f**, Enriched GO categories among significantly downregulated genes in cells of the mesodermal clusters in the mutant embryos ordered by false discovery rate (FDR). **g**, STRING network of all significantly downregulated genes in cells of the mesodermal clusters in the mutant embryos. Nodes not connected to the main network have been removed. Nodes belonging to GO category “embryonic development” colored in blue; nodes belonging to the STRING network cluster “Wnt signaling pathway, and TGF-beta signaling pathway” colored in red. **h**, volcano plot showing the DEGs between amnion (AM-1 and AM-2) of the wildtype and mutant embryos. Grey areas show expected group mean difference from a random cell subset and a false discovery rate (FDR) of 1%. Red labelling indicates that the gene is part of the ISL1 regulon identified by the gene regulatory network interference. **i**, violin plot of the expression levels of *BMP4* and *WNT6* in the different cell types identified in (**c**) separated between wildtype (blue) and mutant (red) embryos. Epi, epiblast; Endo, endoderm; ExE-Meso, extraembryonic mesoderm; ExE-Mech, extraembryonic mesenchyme; AM-1, amnion 1; AM-2, amnion 2; meso-1, mesoderm 1; meso-2, mesoderm 2; wt, wildtype; mt, mutant.

Integration of this dataset with scRNA sequencing data of an *in vivo* human Carnegie stage 7 embryo ^36^ showed, that the mesodermal cell populations underrepresented in the mutant embryos match to the primitive streak and nascent mesoderm clusters of the human gastrula (Fig. 3e). This strongly supports the conclusion that mutant primate embryos fail to form mesoderm.

### Loss of *ISL1* in amnion impairs signaling

Differential gene expression analysis between the cells in the mesodermal clusters (mesoderm 1 and mesoderm 2) of mutant and wildtype embryos revealed 251 genes that were significantly downregulated in the mutants. This list was enriched for GO terms (biological processes) such as “anterior/posterior pattern specification”, “embryo development” and “mesoderm development” (Fig. 3f). 163 of these genes were members of a large, highly interconnected STRING network (PPI enrichment p-value: < 1.0e-16). Within this network, a STRING network cluster termed “Wnt signaling pathway, and TGF-beta signaling pathway” showed the highest enrichment and was located in the center, suggesting that alterations in Wnt and/or TGF-beta signaling could be underlying the observed phenotype (Fig. 3g).

The amnion has previously been suggested to serve as a signaling hub for gastrulation ^37^ and, due to its high expression of *ISL1*, is likely to be the primary affected tissue in the mutant embryos. Differential gene expression analysis in amniotic cells of mutant versus wildtype embryos revealed 184 significantly downregulated genes in the mutant amnion, of which a significant proportion were members of the identified ISL1 regulon (Fig. 3h-i). Among those was *BMP4*, a secreted member of the TGF beta signaling pathway and a known downstream target of ISL1 in mice ^38^, as well as *WNT6*, a secreted Wnt-ligand (Fig. 3h). BMP4 was previously shown to be essential for murine mesoderm formation ^39^ as well as for inducing primitive streak like cells from hESCs ^34^, while WNT6 is known to be required later in embryonic development in somatogenesis in mouse and chick ^40, 41^.

The formation of primordial germ cells (PGCs) in primate embryos has previously been suggested to occur in the amnion in response to BMP4-signaling ^42^. We identified PGCs in our dataset as outliers (> 4 times standard deviation) of a PGC-specific *(DAZL, DPPA3, SOX17, PRDM1, TFAP2C, NANOS3, NANOG)* gene signature score (Supplementary Fig. 4e-f). PGCs were located within the mesodermal cell clusters of wildtype and mutant embryos (Supplementary Fig. 4g-h), suggesting that the loss of *ISL1* is not interfering with PGC formation.

Taken together, these findings suggest that loss of *ISL1* causes altered signaling from the amnion, which results in failure to form mesoderm in the early NHP embryo eventually leading to embryonic lethality.

### *ISL1*-null hESC-derived amnion fails to induce mesoderm

To validate this conclusion and to investigate whether these findings also apply to humans, *ISL1-*null hESCs (*ISL1*-null) harboring the most abundant long deletion in the *ISL1* locus found in the mutant embryos (Supplementary Fig. 1a and 5a) were generated and analyzed *in vitro* (Fig. 4a). Amniotic ectoderm-like cells (AMLCs) derived from the *ISL1*-null showed a 50% reduction in *ISL1* mRNA (Supplementary Fig. 5b) and absence of ISL1 protein which was abundantly expressed in wildtype hESCs derived AMLCs (Fig. 4b). We noticed a slight, nonsignificant reduction in *WNT6* and a significant reduction in *BMP4* expression of about 50%, confirming the functional defect in *ISL1*-null derived AMLCs (Fig. 4c, Supplementary Fig. 5c). In line with the findings from the *in vitro* cultured embryos AMLCs derived from the *ISL1*-null failed to induce mesoderm-like cells (MeLCs) from hESCs shown by a strong reduction in the number of Brachyury (BRA, *TBXT)* expressing cells in the lower compartment (Fig. 4d-e). Notably, *TBXT* expression levels were similar between *ISL1*-null and wildtype using a directed differentiation protocol towards mesoderm-like cells ^34^ (Supplementary Fig. 5d), highlighting that the failure to form mesoderm in the *ISL1*-null is a non-cell autonomous defect caused by altered signaling from AMLCs.

**Fig. 4:**
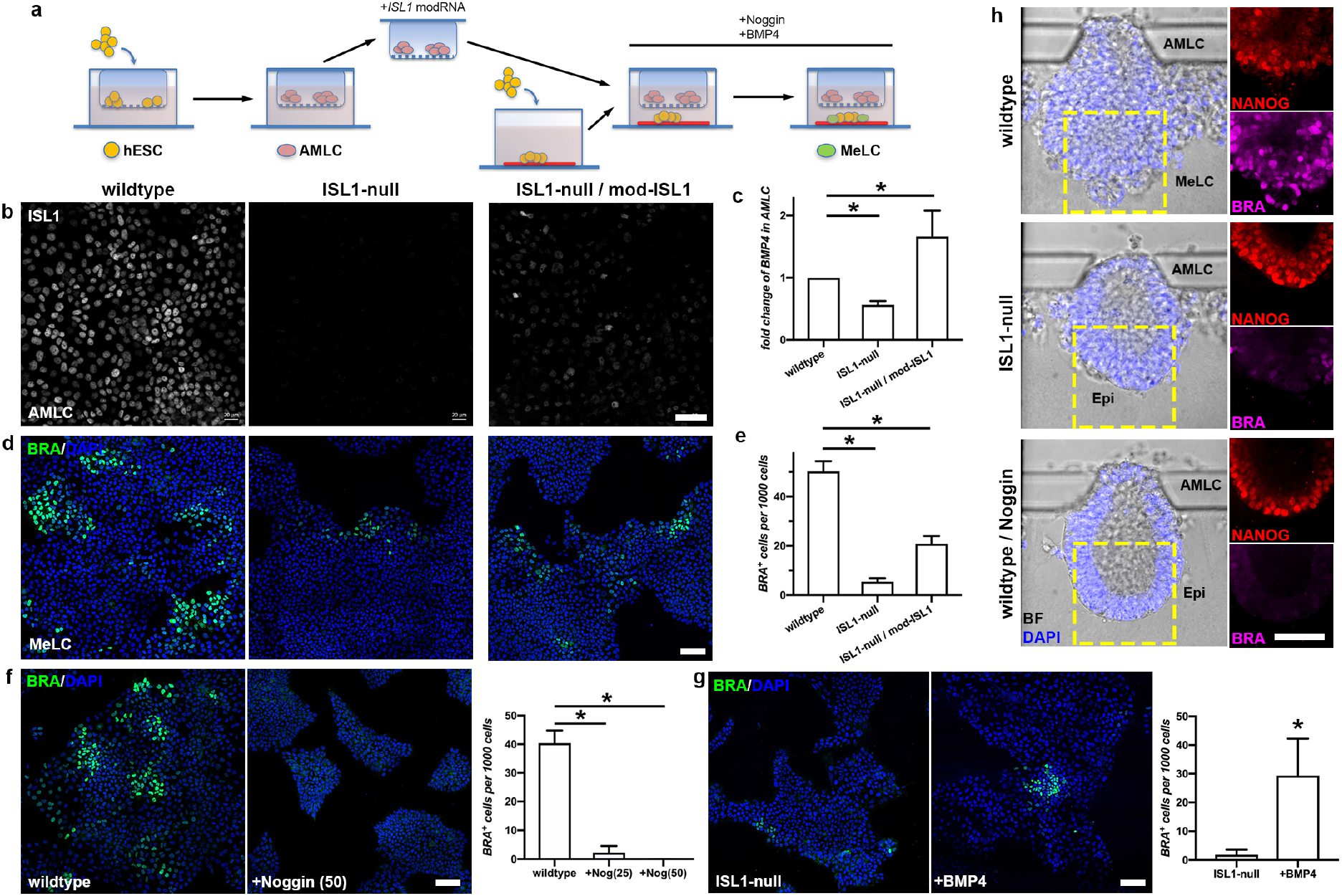
*ISL1* regulates human mesodermal cell formation through BMP4 pathway. See also Supplementary Fig. 5 and 6. **a**, a diagram of the transwell assay. **b**, ISL1 protein in AMLCs. **c**, the expression of *BMP4* in AMLCs. **d**, mesoderm marker Brachyury (BRA, green) in MeLCs. Nucleus shown by DAPI (blue). **e**, quantification of BRA+ cells from (**d**). **f**, Brachyury signal in wildtype MeLC treated with or without Noggin, followed by quantification of BRA+ cells. **g**, Brachyury signals in *ISL1*-null MeLC treated with or without BMP4, followed by quantification of BRA+ cells. **h**, the morphology (left panel) of embryonic-like sacs overlayed with nucleus shown by DAPI (blue). NANOG (red) protein and BRA (magenta) protein on right panels. n = 30 for each. Scale bar, 50 μm. Analyzed by student’s t-test, n=4 for each condition. Data are represented as Mean ± SEM, significance levels are depicted by asterisks: * p-value <0.05.

Modified mRNA-based re-expression of *ISL1* in *ISL1*-null AMLCs restored *BMP4* expression confirming that BMP4 is acting downstream of ISL1 (Fig. 4c). Moreover, it significantly increased the capacity of the *ISL1*-null AMLCs to induce BRA positive cells (Fig. 4d-e).

### BMP4 rescues mesoderm formation *in vitro*

To further investigate whether the formation of MeLCs is indeed depending on BMP4-signaling from the AMLCs, we inhibited BMP4 downstream signaling by using Noggin. This resulted in a dose-dependent reduction in the number of BRA positive cells in wildtype hESCs (Fig. 4f), mimicking the *ISL1*-null phenotype. Reversely, external addition of BMP4 in the *ISL1*-null led to a partial rescue of the phenotype, shown by a significant increase in the number of BRA expressing cells (Fig. 4g), suggesting that BMP4-signaling is responsible in part for the observed phenotype.

The capacity of *ISL1-*null and wildtype hESCs to self-organize into an embryonic-like sac was assessed in a microfluidic system that has been shown to faithfully recapitulate the peri-implantation development of the epiblast lineages ^37^. We noticed that the high-dose of BMP4 (50 ng/ml) used in the protocol of the original publication masked the phenotype in the *ISL1*-null and thus, we reduced the BMP4 concentration (25 ng/ml). With this reduced BMP4 dose, the wildtype cells still showed proper formation of embryonic-like sacs, adequate break of symmetry and formation of MeLCs in the epiblast-like region as shown by positive staining for BRA (Fig. 4h and Supplementary Fig. 5e). In these conditions, *ISL1-*null cells were capable of self-organizing into embryonic-like sacs and broke symmetry similar to the wildtype but failed to develop further. The epiblast-like cells remained in a columnar shape with high levels of the pluripotency factor NANOG ^24^ and stained negative for BRA (Fig. 4h and Supplementary Fig. 5e), indicating failure to form MeLCs similar to the findings observed in the mutant embryos during *in vitro* culture. A similar phenotype was observed in embryonic-like sacs from wildtype hESCs when BMP4 downstream signaling was inhibited by using high-dose Noggin highlighting the importance of BMP4 signaling for MeLCs formation (Fig. 4h and Supplementary Fig. 5f).

## Discussion

In this study we generated a high-resolution developmental roadmap of post-implantation NHP embryos and identified the amnion as a key signaling structure essential for mesoderm formation in primates. NHP embryos hypomorphic for the transcription factor *ISL1*, which is highly expressed in primate amnion, failed to form mesoderm due to a reduction in BMP4-signaling and are not capable of giving rise to viable offspring (Fig. 5).

**Fig. 5:**
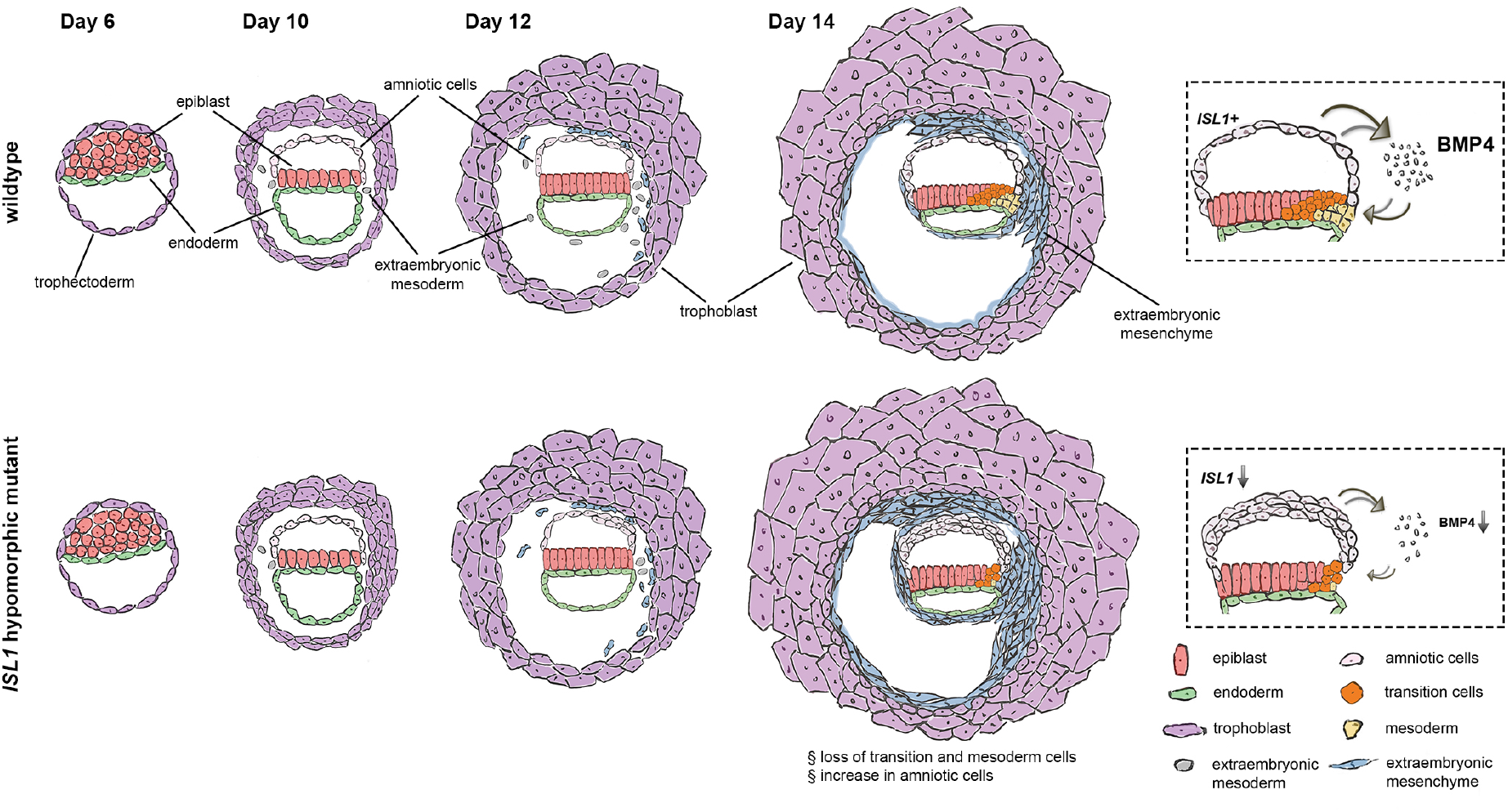
A summary scheme depicting the embryogenesis of wildtype and *ISL1* hypomorphic mutant embryos.

Notably, the role of ISL1 acting upstream of BMP4 seems to be a conserved pathway. The complete loss of Isl1 in mice leads to normal gastrulation since, it is not expressed in the mouse embryo before E7.0 ^7, 32^. However, it is embryonic lethal at approximately E10.5 due to severe cardiac defects accompanied by a strong reduction in *Bmp4* ^7^, suggesting that BMP4 is acting downstream of ISL1 in cardiac development. Similar observations linking *Isl1* and *Bmp4* were also made in mice during genital development ^43^ and embryonic limb formation ^44^.

It is known that the initiation of mesoderm formation is dependent on BMP4 signaling, which is provided by the extraembryonic ectoderm in mice ^11, 45^. The findings from our study suggest that this role is taken over by the amnion during primate embryogenesis. Mouse and primate embryos have a similar appearance before the implantation stage, although the transcriptome already differs in key aspects ^10, 46^. After implantation, the structural differences between mouse and primate embryos become more evident. Mouse embryos form a cup-like structure, while primate embryos acquire a disk-like shape and have a prominent amnion, which is absent in mouse embryos before gastrulation ^28, 47^. Although the signaling network guiding gastrulation appears to be largely conserved across species ^36^, the findings from our study show that the anatomical differences in the early embryos are associated with the presence of alternative signaling centers.

Explanations for differences in gene essentiality between humans and mice span two ends of a spectrum. When an essential gene in mice fails to demonstrate a similar phenotype in humans the disparate human phenotype could either be very subtle, or the opposite, especially severe. We initially hypothesized that the low frequency of damaging *ISL1* variants reported in human cohorts was due to an important role in early embryonic development. Indeed, we detected its requirement during gastrulation. It is possible that comparable effects could also be observed for other genes, with a similar discrepancy between mouse and human phenotypes. This study highlights that *in vitro* cultured primate embryos are a powerful tool to model key steps of early human development and could make an important contribution in addressing these questions.

Further advances in the *in vitro* culture systems might enable us to support the embryo longer and to study early organogenesis, including the emergence of cardiac progenitor cells in lateral plate mesoderm. This would enrich our knowledge on human embryogenesis, help to identify causes for pregnancy loss and congenital malformations and, eventually, open the avenue for new therapies.

## Methods

### Cynomolgus macaque

Healthy cynomolgus monkeys (Macaca fascicularis), ranging from 5 to 12 years old, were used in this study. All animals were housed either at the facility of Yunnan Key Laboratory of Primate Biomedical Research in China, or at Astrid Fagræus laboratory in Karolinska Institutet in Sweden. Both facilities are accredited by AAALAC international. The ethics and all experimental protocols were approved in advance by the Institutional Animal Care and Use Committee of LPBR in China (KBI K001115033/01,01) and by the Jordbruksverket in Sweden (ethical permit number N277/14). Animals involved in this study were never used for other treatments.

### *In vitro* fertilization

NHP embryos were collected as described in previously publication ^48^. In brief, healthy female monkeys aged 5-8 years with regular menstrual cycles were selected as oocyte donors. Before superovulation, female animals were treated with rhFSH for 8 days, and administrated rhCG on day 9. Oocytes were collected by laparoscopic follicular aspiration 32-35 hours after rhCG administration. MII (first polar body present) oocytes were performed with intracytoplasmic sperm injection to generate zygotes and the fertilization was confirmed by the presence of two pronuclei. Zygotes were cultured in embryo culture medium-9 (HECM-9) containing 10% fetal calf serum in 37°C incubator supplying 5% CO2 until blastocyst stage. Blastocysts were then used for embryo transfer or post-implantation *in vitro* culture. The homemade HECM-9 contains polyvinyl alcohol (0.1 mg/mL), calcium chloride (1.9 mM), magnesium chloride (0.46 mM), potassium chloride (3.0 mM), sodium chloride (113.8 mM), sodium bicarbonate (25.0 mM), sodium lactate (4.5 mM), MEM amino acid, MEM non-essential amino acid, and Gentamicin (10 mg/mL).

### Embryo transfer and pregnancy diagnosis

Embryos were transferred into the oviducts of the matched recipient monkey as described in previous study ^49^. A total of 27 female monkey recipients with proper hormone level of β-estradiol and progesterone were used as surrogate recipients. Each recipient received 2-4 blastocysts. The pregnancy was primarily diagnosed by ultrasonography at 2-3 weeks after embryo transfer. Clinical pregnancy and the number of fetuses were confirmed by fetal cardiac activity and the presence of gestation sacs. When terminating pregnancy, caesarean section was performed. Tissue from umbilical cord, ear and tail was collected for genotyping.

### Generation of *ISL1* hypomorphic mutant embryos

NHP zygotes were injected with the mix of Cas9 protein and guide RNAs. Intracytoplasmic injections were performed with a Nikon microinjection system under standard conditions. The embryos were cultured in HECM-9 supplemented with 10% fetal calf serum in 37°C incubator supplying 5% CO2. Genetic modified embryos with high quality from morula to blastocyst stage were used for further studies.

### *In vitro* embryo culture

To culture blastocyst beyond implantation stage, we applied an optimized protocol based on the human embryo culture protocol from Zernicka-Goetz’s group ^50^. Frozen NHP blastocysts were thawed right before culturing by using the Thawing Media (Kizatato) and cultured in blastocyst culture medium (Origio), for at least 4 hours to recover. Blastocysts were then treated with Acidic Tyrode’s solution to remove the Zona pellucida and transferred to an ibiTreat 8-well μ-plate (Ibidi) containing 300 μL of pre-equilibrated in vitro culture medium 1 (IVC1). On the second day, 150 μL of IVC1 was carefully removed and 200 μL pre-equilibrated in vitro culture medium 2 was added. Blastocyst growth was monitored and medium was changed every two days until termination of experiments.

### Single cell dissociation and RNA sequencing

NHP embryos at Day 10, Day 12 and Day 14 were washed with PBS and treated with TrypLE Express Enzyme for 30 minutes at 37°C. After incubation, samples were gently dissociated into single cells by mouth pipetting. Single cells were transferred into a RNase-free, low-cell-adhesion 1.5 mL tube and centrifuged at 300 g for 5 minutes. Supernatant, containing some remaining cells was transferred into a new tube for genotyping. The cell pellet was resuspended with 40 μL of PBS containing 2% Bovine Serum Albumin. Cells were loaded into the 10x Genomics Chromium system within 30 minutes after dissociation. 10x Genomics v3 libraries were prepared according to the manufacturer’s instructions. Libraries were sequenced with a minimum coverage of 30,000 raw reads per cell on an Illumina NovaSeq with paired-end sequencing.

### Reads mapping, gene expression counting and correction

Sequencing data was aligned and quantified by using the Cell Ranger Pipeline v3.1.0 against the ensemble genome build Macaca_fascicularis_5.0 release 96 ^51^. Ambient RNA contamination was estimated through the levels of choriogonadotropins expression in epiblast (POU5F1 positive) cells and removed from the count matrix using SoupX ^52^. A gene was retained for analysis if it showed expression in at least 3 cells. Each sample was filtered based on expression level of mitochondrial genes (below 7.5%) and number of expressed genes. Details on the estimated contamination in each sample, the filtering criteria and number of cells retained for the analysis are provided in Supplementary Fig. 2a.

### Reads mapping and gene expression counting of *in vivo* dataset

The raw, archived single cell RNA sequencing data from in vivo cynomolgus embryos ^19^ was downloaded from the GEO database (GSE74767) and processed using TrimGalore v0.6.1. The reads passing quality control were aligned against the ensemble genome build macaca_fascicularis_5.0 release 96 using STAR v2.5.3 and counted using featureCounts v1.5.2. Cells expressing at least 1000 genes were kept for the integration with our dataset.

### Data integration, dimensionality reduction and clustering

For analysis of the single cell RNA sequencing data from the wildtype in vitro cultured embryos, we integrated the filtered, corrected count matrices of the different batches for each day separately using the reciprocal principal component analysis (PCA) approach implemented in the Seurat package v3.1.3 ^53, 54^ based on 30 dimensions and 5000 anchor features. After integration, we performed PCA analysis on the integrated data followed by embedding into low dimensional space with Uniform Manifold Approximation and Projection (UMAP) as implemented in the R-package ‘uwot’. For clustering, the shared Nearest Neighbor (SNN) Graph was constructed on the UMAP embedding by calling the FindNeighbors() function followed by the identification of clusters using the FindClusters() function, both part of the Seurat package. In some samples this clustering approach separated large, homogeneous cell groups into small sub-clusters with no distinct biological meaning. In these cases, the clusters were re-combined manually and both, the unsupervised and the manually adjusted clustering, was reported in the manuscript.

To integrated the single cell RNA sequencing data from in vivo cynomolgus embryos ^19^ with our dataset we combined the three time points (Day 10, Day 12 and Day 14) from our dataset for each batch separately and did the same for the in vivo data from embryonic day 8 (E08), E09, E13 and E14 resulting in three separate datasets. Normalization, Scaling and PCA was performed separately on each of these datasets after which they were combined using the reciprocal PCA approach described above based on 30 dimensions and 2000 anchor features.

For the analysis of the wildtype and *ISL1* hypomorphic mutant embryos the different batches were integrated separately for each day by the same reciprocal PCA approach outlined above based on 30 dimensions and 5000 anchor features using the wildtype datasets as reference. Dimensionality reduction and clustering was performed as described above.

To integrated our dataset with the single cell RNA sequencing data from the in the vivo human Carnegie stage 7 embryo ^36^, Macaca fascicularis gene IDs were converted to homo sapiens gene symbols using the according orthologue list from Ensemble. Normalization, Scaling and PCA was performed separately on each of the batches (wildtype and mutant embryos Day 14) from our dataset and the human dataset separately after which they were combined using the reciprocal PCA approach described above based on 30 dimensions and 2000 anchor features. Dimensionality reduction and clustering was performed as described above.

### Differential gene expression analysis

Mainly due to the differences in cell numbers we observed a significant variation in sequencing depth between samples in our dataset (Supplementary Fig. 2). It has recently been shown, that the effect of differences in read depth on differential gene expression analysis can be minimized by using regularized negative binomial regression as implemented in the R-package SCtransform ^55^. Thus, all differential gene expression analysis was performed using a t-test on Pearson residuals after SCtransformation of the raw, filtered counts of the integrated Seurat object as implemented previously ^55^. Gene expression data depicted throughout the manuscript in feature plots or violin plots are SCtransformed data. Expression data depicted in heatmaps are scaled, log-transformed expression values normalized to the total counts for each cell calculated through running the NormalizeData() function followed by the ScaleData() function from the Seurat package.

For analysis of protein-protein interactions and enriched GO terms among differentially expressed genes, Macaca fascicularis gene IDs were converted to homo sapiens gene symbols using the according orthologue list from Ensemble. Interaction networks of differentially expressed genes were created using STRING v11.0 and analyzed for enriched GO terms as well as enriched STRING network clusters using standard settings ^56^.

### Visualization of gene signatures

Scoring and visualization of gene signatures was performed using the Single Cell Signature Explorer v3.1 ^57^. Gene signatures were created by identifying orthologues for the genes that have been previously described to mark the naïve and primed state of pluripotency in human embryonic stem cells ^21^ in the Macaca fascicularis genome using the according orthologue list provided from Ensemble through BioMart ^58^.

### Pseudotime analysis

Pseudotime analysis was performed using Monocle3 v0.2 ^59–61^. The principal graph was learned on the UMAP embedding extracted from the integrated Seurat object. Differentially expressed genes were calculated on the raw, filtered count matrix extracted from the integrated Seurat object using the Moran’s I test implemented in the graph_test() function from the Monocle3 package. The genes were ranked according to their Moran’s I and the top 100 genes were selected for display in the heatmap.

### Gene regulatory network analysis

Gene regulatory network (GRN) analysis was performed using the R-package SCENIC (Single Cell rEgulatory Network Inference and Clustering) v1.1.2-2 ^31^ and the command line interface (CLI) of the python implementation pySCENIC. Macaca fascicularis gene IDs were converted to homo sapiens gene symbols using the according orthologue list from Ensemble. The raw, filtered count matrix extracted from the integrated Seurat object was pre-filtered and genes with at least 39 counts, equal to at least 3 UMI counts in 1% of the cells, present in at least 13 cells, equal to 1% of the cells, were used as input for the CLI. The human motif collection v9 and the cisTarget databases for hg38 were used in the pipeline and downloaded from https://resources.aertslab.org/cistarget/. Thresholds used for binarization were derived from the AUC values using Hartigan’s Dip Test (HDT). After binarization, regulons showing activity in at least 1% of the cells were included in the downstream analysis.

### Culture of human embryonic stem cells

Human embryonic stem cells used in this study include HES-3 human ES cells and H9 human ES cells (WiCell). Genetic modification of the *ISL1* locus to generate *ISL1-null* hESCs was performed on HES-3 cells by applying CRISPR/Cas9 with the same guide RNAs used in NHP blastocysts. Two *ISL1* knockout cell lines were generated, named ISL1_ko_c15 and ISL1_ko_c51, and genotyped (Supplementary Fig. 5a). All cell lines were authenticated as karyotypically normal by Cell Guidance Systems (United Kingdom) (Supplementary Fig. 5a). Mycoplasma contamination test was performed regularly as negative. hESCs were maintained in a standard feeder-free culture system using mTeSR1 medium on 1% Matrigel or Essential 8 medium on 1% Vitronectin. Cells were passaged every 4-5 days and visually examined during each passage to ensure absence of spontaneously differentiation. Work with human embryonic stem cells was carried out according to Swedish legislation following the recommendations of the Swedish National Council on Medical Ethics.

### Genotyping

Genomic DNA was extracted by Phenol-Chloroform method. DNA fragment covering both guide RNA target sites were PCR amplified and ligated to TOPO TA cloning vector. At least 50 bacteria clones per sample were picked for Sanger sequencing and used to estimate the genomic mutation rate. The transcriptomic mutation rate of *ISL1* hypomorphic mutants was also calculated. cDNA libraries of each scRNA-sequencing sample were used to amplify the *ISL1* mRNA fragment covering both guide RNA target sites. PCR products were ligated into TOPO TA cloning vector. At least 50 clones per cDNA library sample were picked and performed Sanger sequencing.

### Off-target assay

Cas-OFFinder was applied to search for potential off-target sites with maximal two mismatches and two bulges ^18^. Among all off-target candidates of both gRNAs, targets located on gene exons were selected for test. The DNA fragments of target sites were PCR amplified and the sequences were confirmed by Sanger sequencing.

### RNA extraction and quantitative real-time PCR

Total message RNA was extracted by Direct-zol RNA miniprep kits and reverse transcription to cDNA library was prepared by GoScript Reverse Transcriptase. Quantitative real-time PCR was performed by PowerUp SYBR Green Master Mix on ABI 7500Fast machine.

### Transwell assay

The transwell assay was performed based on previous work by Zheng and colleagues ^37^. In brief, it was performed on Transwell 12-well plates with permeable polyester membrane inserts (0.4 μm, Corning). The membrane inserts were coated with 1% Geltrex diluted in DMEM/F12 for 1 hour before use. hESCs were collected and re-suspended in culture medium containing Y-27632 (10 μM) and seeded onto the membrane insert at a density of 3 x 10^4^ cells per cm^2^. Eighteen hours after seeding, culture medium was changed to E6 medium supplemented with bFGF (20 ng/mL) and BMP4 (50 ng/mL) and cultured for 48 hours. On day 3, undifferentiated hESCs were collected, re-suspended in E6 supplemented with bFGF (20 ng/mL) and seeded at a density of 9 x 10^4^ per well on freshly coated 12-well plates. The membrane inserts were washed with E6 + bFGF and transferred on top of the re-seeded hESCs. Cells were collected after 48 hours for analysis. Two wildtype hESC-lines (HES-3 and H9) and two *ISL1-*null lines were used in this assay. Both of the wildtype cell lines showed comparable results, as did the two *ISL1-* null lines.

Noggin inhibition, BMP4 rescue and *ISL1* modified mRNA rescue were performed on transwell assay as well. As shown in Fig. 4a, Noggin (50 ng/mL) or BMP4 (20 ng/mL) were administrated into E6+bFGF in the lower part of the transwell inserts from day 3, after transferring inserts on top of hESCs. The *ISL1* modRNA was designed and *in vitro* synthesized according to the previous work ^62^. 1mg of purified *ISL1* modRNA was introduced into each sample of amniotic like cells on insert membrane on day 3 and then transferred the inserts on top of hESCs to induce the mesodermal like cell formation.

### Primitive streak induction from hESCs

Differentiation of hESCs to primitive streak-like cells was done in chemically defined media as previously described ^34^. In brief, posterior primitive streak was induced by the addition of bFGF (20 ng/ml), the phosphoinositide 3-kinases (PI3K)-inhibitor LY294002 (10 μM) and BMP4 (10 ng/ml). Anterior primitive streak was induced with the same factors and, additionally, Activin A (50 ng/ml). After 40 hours cells were harvested. RNA extraction, reverse transcription and quantitative real-time PCR were performed as detailed below with 200 ng RNA as input for RT-reaction. All experiments were performed in at least biological triplicates.

### Microfluidic assay of embryonic-like sac

This assay was performed as previously described ^63^. Briefly, the microfluidic device is fabricated by bonding a PDMS structure layer to a coverslip. Geltrex is diluted to 70% using E6 medium and loaded into the central gel channel separated from the side channels by trapezoid-shaped supporting posts. Upon gelation, Geltrex matrix would generate concave Geltrex pockets between supporting posts for cell seeding. hESCs suspended in mTeSR1 medium was introduced into the cell loading channel and allowed to settle and cluster in the gel pockets. After hESCs cluster formation, mTeSR1 medium was replaced by a basal medium (E6 and 20 ng/mL bFGF), and 20 ng/mL BMP4 was supplemented only into the cell seeding channel. After 18 hours of BMP4 stimulation, the BMP4 medium was replaced by the basal medium. The microfluidic devices were fixed at 48 hours since the hESCs clusters were exposed to BMP4. To test the BMP4 signaling function, Noggin (50 ng/mL and 500 ng/mL) was supplemented into the basal medium into the cell loading channel for 48 hours. The hESCs clusters were then fixed and stained.

### Cryosection of NHP embryos

Day 14 NHP embryos were fixed by 2% paraformaldehyde overnight at 4°C and then washed by PBS. Dehydrate the fixed embryos by 30% sucrose overnight at 4°C, and then embedded in OCT and froze in liquid nitrogen. Frozen blocks were performed cryosection on Cryostat (Thermo CryoStar NX70) according to manufacturer’s protocol.

### Immunohistochemistry

Immunohistochemistry of cells from the transwell assay was performed following standard procedures. Briefly, cells were fixed in 2% paraformaldehyde for 30 minutes at room temperature and washed with PBS. Cells were blocked in blocking buffer (serum diluted in PBS with 0.1% Triton X-100) for one hour and then incubated with primary antibodies diluted in blocking buffer overnight at 4°C. Cells were washed with PBS supplemented with 0.1% Tween-20 (PBS-T) and incubated with secondary antibodies diluted in blocking buffer for 2 hours at room temperature. After incubation, secondary antibodies were washed off by PBS-T, and the samples were mounted for imaging. Staining of embryonic-like sac structure was performed as previously described ^63^. Antibodies used are listed in the Key Resources Table. Confocal micrographs were acquired by Zeiss 700 LSM confocal microscope or Olympus spinning-disc confocal microscope (DSUIX18) equipped with an EMCCD camera (iXon X3, Andor). The bright-field morphologic images of embryonic-like sacs were acquired by Zeiss Observer.Z1 microscope equipped with a monochrome CCD camera (AxioCam, Carl Zeiss MicroImaging). Images were analyzed by iMaris.

### Quantification and Statistical Analysis

Values are shown as the mean value plus SEM. Continuous data was analyzed using student’s t-test. P-values or adjusted p-values (where appropriate) below 0.05 were considered statistically significant. Details on the samples (e.g. number of biological replicates) are indicated in figure legends. Graphs were generated using Prism or R.

## Data and Code Availability

The raw data, unfiltered count matrix and processed count matrix are deposited in the Gene Expression Omnibus (GEO) database with the accession number GSE148683 and will be publicly released upon publication. All code is available from the authors upon request.

## Acknowledgments

We thank the Eukaryotic Single Cell Genomics facility, hosted at SciLifeLab, for single cell RNA sequencing service. The computations were performed on resources provided by SNIC through Uppsala Multidisciplinary Center for Advanced Computational Science (UPPMAX) under Project SNIC 2019/8-234. Martin Dahlö at UPPMAX is acknowledged for assistance concerning technical and implementational aspects in making the code run on the UPPMAX resources. We would like to thank Xiaobing He and Jesper Sohmer for technical support and want to acknowledge Dr. Xidan Li for advice on the bioinformatic analysis. This study was supported by grants from the National Key Research and Development Program (2016YFA0101401) and the Major Basic Research of Yunnan (2019FY002) from China, and a Grant for Swedish-Chinese collaboration from the Swedish Research Council (Dnr 539-2013-7002). K.C. was supported by a Grant for distinguished professors of the Swedish Research Council (Dnr 541-2013-8351). F.L. was supported by Ming Wai Lau Centre for Reparative Medicine, Knut and Alice Wallenberg Foundation, Centre for Innovative Medicine, Swedish Research Council, Ragnar Söderberg Foundation and Wallenberg Academy. A.G. was supported by a grant from the German research foundation (grant number: GO3220/1-1). Y.Z. and J.F. were supported by the National Institutes of Health (R21 NS113518) and the National Science Foundation (CMMI 1917304 and CBET 1901718).

## Author Contributions

K.C. and W.J established the initial joint project and the specific research plan was conceived by K.C. and R.Y.. R.Y., A.G., F.L. and K.C. planned experiments, analyzed data interpreted the results and wrote the manuscript. C.L. and R.Y. designed the guide RNAs. C.C. tested the guide RNA efficiency on cynomolgus cells and embryos. Y.K. and C.S. generated the wildtype and mutant NHP blastocysts. Z.C. and Y.C. performed the NHP embryo collection and transfer. C.Z thawed the NHP blastocysts. R.Y. performed the *in vitro* culture of embryos, collected samples for scRNA sequencing, did genotyping and off-target assay. A.G. analyzed the scRNA sequencing data, performed the primitive streak induction and related quantitative real-time PCR. Y.X. maintained human cell lines, prepared samples for karyotyping and assisted in many experiments. R.Y. did the transwell assay and related immunofluorescent staining and PCR amplification. Y.Z. and J.F. performed the microfluidic assay and related immunofluorescent staining. P.G. analyzed the genetic *ISL1* variants in the human population. N.W. designed and prepared the modified mRNA. Y.N., W.J., F.L. and K.C. supervised the study. R.Y., A.G. and Y.K. contributed equally to the study. Correspondence can be addressed to K.C., F.L., Y.N., or W.J..

## References

1. Domian, I.J. et al. Generation of functional ventricular heart muscle from mouse ventricular progenitor cells. Science 326, 426–429 (2009).

2. Laugwitz, K.L. et al. Postnatal isl1+ cardioblasts enter fully differentiated cardiomyocyte lineages. Nature 433, 647–653 (2005).

3. Moretti, A. et al. Multipotent embryonic isl1+ progenitor cells lead to cardiac, smooth muscle, and endothelial cell diversification. Cell 127, 1151–1165 (2006).

4. Bu, L. et al. Human ISL1 heart progenitors generate diverse multipotent cardiovascular cell lineages. Nature 460, 113–117 (2009).

5. Sahara, M. et al. Population and Single-Cell Analysis of Human Cardiogenesis Reveals Unique LGR5 Ventricular Progenitors in Embryonic Outflow Tract. Dev Cell 48, 475–490 e477 (2019).

6. Pfaff, S.L., Mendelsohn, M., Stewart, C.L., Edlund, T. & Jessell, T.M. Requirement for LIM homeobox gene Isl1 in motor neuron generation reveals a motor neuron-dependent step in interneuron differentiation. Cell 84, 309–320 (1996).

7. Cai, C.L. et al. Isl1 identifies a cardiac progenitor population that proliferates prior to differentiation and contributes a majority of cells to the heart. Dev Cell 5, 877–889 (2003).

8. Zaidi, S. et al. De novo mutations in histone-modifying genes in congenital heart disease. Nature 498, 220–223 (2013).

9. Jin, S.C. et al. Contribution of rare inherited and de novo variants in 2,871 congenital heart disease probands. Nat Genet 49, 1593–1601 (2017).

10. Petropoulos, S. et al. Single-Cell RNA-Seq Reveals Lineage and X Chromosome Dynamics in Human Preimplantation Embryos. Cell 165, 1012–1026 (2016).

11. Tam, P.P. & Loebel, D.A. Gene function in mouse embryogenesis: get set for gastrulation. Nat Rev Genet 8, 368–381 (2007).

12. Wamaitha, S.E. & Niakan, K.K. Human Pre-gastrulation Development. Curr Top Dev Biol 128, 295–338 (2018).

13. Kessler, D.S. & Melton, D.A. Vertebrate embryonic induction: mesodermal and neural patterning. Science 266, 596–604 (1994).

14. Rossant, J. Genetic Control of Early Cell Lineages in the Mammalian Embryo. Annu Rev Genet 52, 185–201 (2018).

15. Ma, H. et al. In vitro culture of cynomolgus monkey embryos beyond early gastrulation. Science 366, eaax7890 (2019).

16. Niu, Y.Y. et al. Dissecting primate early post-implantation development using longterm in vitro embryo culture. Science 366, 837–+ (2019).

17. Xiang, L. et al. A developmental landscape of 3D-cultured human pre-gastrulation embryos. Nature 577, 537–542 (2020).

18. Bae, S., Park, J. & Kim, J.S. Cas-OFFinder: a fast and versatile algorithm that searches for potential off-target sites of Cas9 RNA-guided endonucleases. Bioinformatics 30, 1473–1475 (2014).

19. Nakamura, T. et al. A developmental coordinate of pluripotency among mice, monkeys and humans. Nature 537, 57–62 (2016).

20. Liu, D. et al. Single-cell RNA-sequencing reveals the existence of naive and primed pluripotency in pre-implantation rhesus monkey embryos. Genome Res 28, 1481–1493 (2018).

21. Messmer, T. et al. Transcriptional Heterogeneity in Naive and Primed Human Pluripotent Stem Cells at Single-Cell Resolution. Cell Rep 26, 815–824 e814 (2019).

22. Shahbazi, M.N. et al. Pluripotent state transitions coordinate morphogenesis in mouse and human embryos. Nature 552, 239–243 (2017).

23. Shakiba, N. et al. CD24 tracks divergent pluripotent states in mouse and human cells. Nat Commun 6, 7329 (2015).

24. Collier, A.J. et al. Comprehensive Cell Surface Protein Profiling Identifies Specific Markers of Human Naive and Primed Pluripotent States. Cell Stem Cell 20, 874–890 e877 (2017).

25. Saykali, B. et al. Distinct mesoderm migration phenotypes in extra-embryonic and embryonic regions of the early mouse embryo. Elife 8, e42434 (2019).

26. Knofler, M. et al. Human placenta and trophoblast development: key molecular mechanisms and model systems. Cell Mol Life Sci 76, 3479–3496 (2019).

27. Rossant, J. & Tam, P.P.L. New Insights into Early Human Development: Lessons for Stem Cell Derivation and Differentiation. Cell Stem Cell 20, 18–28 (2017).

28. Carlson, B.M. Formation of Germ Layers and Early Derivatives, in Human Embryology and Developmental Biology 75–91 (2014).

29. Ross, C. & Boroviak, T.E. Origin and function of the yolk sac in primate embryogenesis. Nat Commun 11, 3760 (2020).

30. Davidson, E. & Levin, M. Gene regulatory networks. Proc Natl Acad Sci U S A 102, 4935 (2005).

31. Aibar, S. et al. SCENIC: single-cell regulatory network inference and clustering. Nat Methods 14, 1083–1086 (2017).

32. Pijuan-Sala, B. et al. A single-cell molecular map of mouse gastrulation and early organogenesis. Nature 566, 490–495 (2019).

33. Chawengsaksophak, K., de Graaff, W., Rossant, J., Deschamps, J. & Beck, F. Cdx2 is essential for axial elongation in mouse development. Proc Natl Acad Sci U S A 101, 7641–7645 (2004).

34. Mendjan, S. et al. NANOG and CDX2 pattern distinct subtypes of human mesoderm during exit from pluripotency. Cell Stem Cell 15, 310–325 (2014).

35. Kitajima, S., Takagi, A., Inoue, T. & Saga, Y. MesP1 and MesP2 are essential for the development of cardiac mesoderm. Development 127, 3215–3226 (2000).

36. Tyser, R.C.V. et al. A spatially resolved single cell atlas of human gastrulation. bioRxiv, 2020.2007.2021.213512 (2020).

37. Zheng, Y. et al. Controlled modelling of human epiblast and amnion development using stem cells. Nature 573, 421–425 (2019).

38. Gao, R. et al. Pioneering function of Isl1 in the epigenetic control of cardiomyocyte cell fate. Cell Res 29, 486–501 (2019).

39. Winnier, G., Blessing, M., Labosky, P.A. & Hogan, B.L. Bone morphogenetic protein-4 is required for mesoderm formation and patterning in the mouse. Genes Dev 9, 2105–2116 (1995).

40. Tajbakhsh, S. et al. Differential activation of Myf5 and MyoD by different Wnts in explants of mouse paraxial mesoderm and the later activation of myogenesis in the absence of Myf5. Development 125, 4155–4162 (1998).

41. Schmidt, C. et al. Wnt 6 regulates the epithelialisation process of the segmental plate mesoderm leading to somite formation. Dev Biol 271, 198–209 (2004).

42. Sasaki, K. et al. The Germ Cell Fate of Cynomolgus Monkeys Is Specified in the Nascent Amnion. Developmental Cell 39, 169–185 (2016).

43. Ching, S.T. et al. Isl1 mediates mesenchymal expansion in the developing external genitalia via regulation of Bmp4, Fgf10 and Wnt5a. Hum Mol Genet 27, 107–119 (2018).

44. Yang, L. et al. Isl1Cre reveals a common Bmp pathway in heart and limb development. Development 133, 1575–1585 (2006).

45. Arnold, S.J. & Robertson, E.J. Making a commitment: cell lineage allocation and axis patterning in the early mouse embryo. Nat Rev Mol Cell Biol 10, 91–103 (2009).

46. Boroviak, T. et al. Single cell transcriptome analysis of human, marmoset and mouse embryos reveals common and divergent features of preimplantation development. Development 145, dev167833 (2018).

47. Pereira, P.N. et al. Amnion formation in the mouse embryo: the single amniochorionic fold model. BMC Dev Biol 11, 48 (2011).

48. Niu, Y. et al. Generation of gene-modified cynomolgus monkey via Cas9/RNA-mediated gene targeting in one-cell embryos. Cell 156, 836–843 (2014).

49. Liu, H. et al. TALEN-mediated gene mutagenesis in rhesus and cynomolgus monkeys. Cell Stem Cell 14, 323–328 (2014).

50. Shahbazi, M., Vuoristo, S., Jedrusik, A., Shahbazi, M. & Zernicka-Goetz, M. Culture of human embryos through implantation stages in vitro. PROTOCOL (Version 1) available at Protocol Exchange (2016).

51. Cunningham, F. et al. Ensembl 2019. Nucleic Acids Res 47, D745–D751 (2019).

52. Young, M.D. & Behjati, S. SoupX removes ambient RNA contamination from droplet based single-cell RNA sequencing data. bioRxiv, 303727 (2020).

53. Butler, A., Hoffman, P., Smibert, P., Papalexi, E. & Satija, R. Integrating single-cell transcriptomic data across different conditions, technologies, and species. Nat Biotechnol 36, 411–420 (2018).

54. Stuart, T. et al. Comprehensive Integration of Single-Cell Data. Cell 177, 1888–1902 e1821 (2019).

55. Hafemeister, C. & Satija, R. Normalization and variance stabilization of single-cell RNA-seq data using regularized negative binomial regression. Genome Biol 20, 296 (2019).

56. Szklarczyk, D. et al. STRING v11: protein-protein association networks with increased coverage, supporting functional discovery in genome-wide experimental datasets. Nucleic Acids Res 47, D607–D613 (2019).

57. Pont, F., Tosolini, M. & Fournié, J.J. Single-Cell Signature Explorer for comprehensive visualization of single cell signatures across scRNA-seq datasets. Nucleic Acids Research 47, e133–e133 (2019).

58. Smedley, D. et al. The BioMart community portal: an innovative alternative to large, centralized data repositories. Nucleic Acids Res 43, W589–598 (2015).

59. Cao, J. et al. The single-cell transcriptional landscape of mammalian organogenesis. Nature 566, 496–502 (2019).

60. Qiu, X. et al. Reversed graph embedding resolves complex single-cell trajectories. Nat Methods 14, 979–982 (2017).

61. Trapnell, C. et al. The dynamics and regulators of cell fate decisions are revealed by pseudotemporal ordering of single cells. Nat Biotechnol 32, 381–386 (2014).

62. Zangi, L. et al. Modified mRNA directs the fate of heart progenitor cells and induces vascular regeneration after myocardial infarction. Nat Biotechnol 31, 898–907 (2013).

63. Zheng, Y. & Fu, J. Protocol for controlled modeling of human epiblast and amnion development using stem cells. PROTOCOL (Version 1) available at Protocol Exchange (2019).

