## Supplementary Figures for "Amnion signals are essential for mesoderm formation in primates"

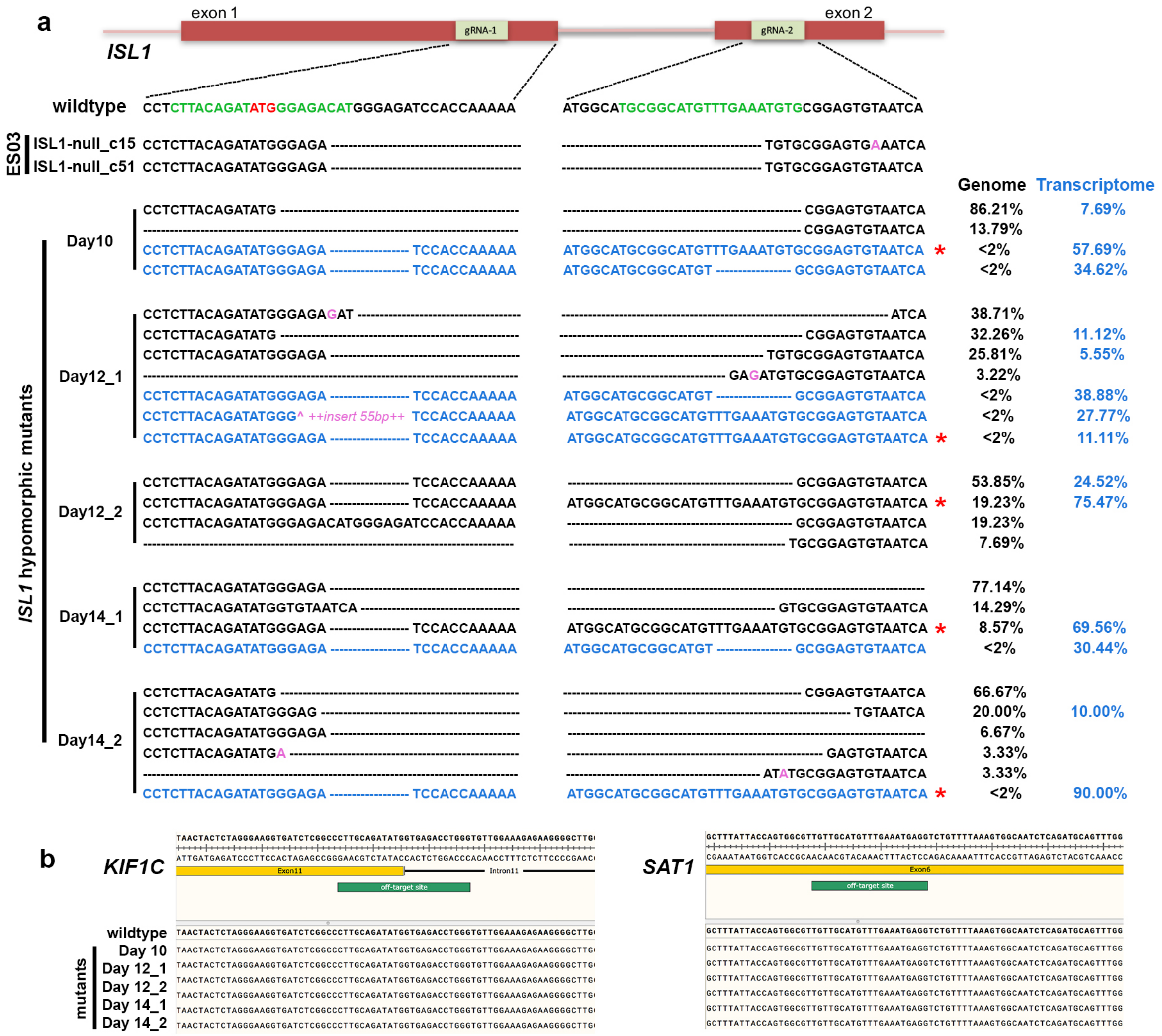


**Supplementary Fig. 1: genotyping and off-target assay of *ISL1* hypomorphic mutants.** Related to Fig. 1, 3 and 4.

**a**, genotyping of two *ISL1*-null hESC-lines and all *ISL1* hypomorphic mutant embryos used for scRNA sequencing. Green sequences mark the gRNA target regions, dotted lines show deletions and pink color indicates insertions. Start codon of *ISL1* is marked in red. Genomic genotyping is shown in black and the genotyping of transcriptome in blue. Numbers are percentages of each mutation type in each sample. Asterisks indicate mutated but functional sequences. **b**, off-target analysis of CRISPR/Cas9 generated *ISL1* hypomorphic mutants.

**a**

| type | day | batch | embryos per batch | cells from cellranger | estimated % contamination | minimal gene # used for filtering | cells after filtering |
|---|---|---|---|---|---|---|---|
| wt | D10 | B1 | 2 | 758 | 3.22 | 2500 | 658 |
| wt | D10 | B2 | 3 | 874 | 0.20 | 2500 | 670 |
| mt | D10 | B1 | 3 | 1063 | 1.04 | 2500 | 769 |
| wt | D12 | B1 | 1 | 510 | 2.27 | 2500 | 325 |
| wt | D12 | B2 | 2 | 1333 | 1.08 | 1500 | 981 |
| mt | D12 | B1 | 3 | 7263 | 4.35 | 1500 | 6018 |
| mt | D12 | B2 | 1 | 8284 | 7.99 | 1500 | 5842 |
| wt | D14 | B1 | 2 | 4685 | 2.12 | 1500 | 3176 |
| wt | D14 | B2 | 2 | 1731 | 1.02 | 1500 | 1384 |
| mt | D14 | B1 | 3 | 12951 | 6.90 | 1000 | 9625 |
| mt | D14 | B2 | 1 | 7719 | 3.22 | 1500 | 3882 |

**b**

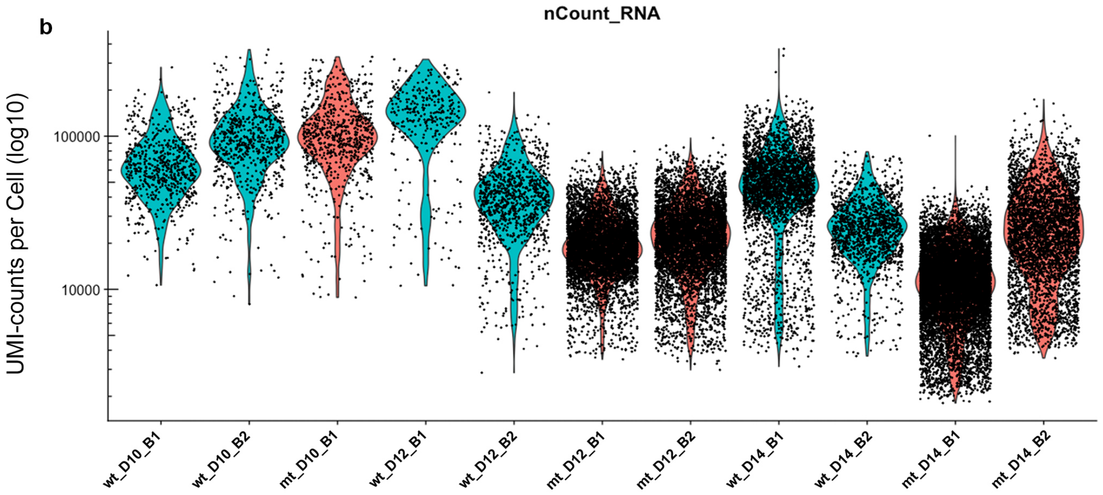


**c**

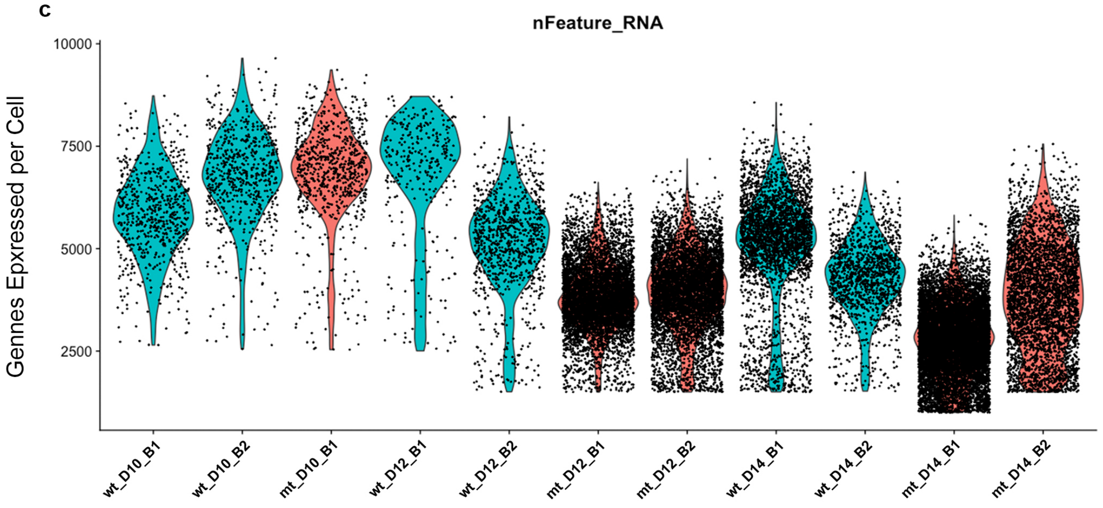


**Supplementary Fig. 2: Cell numbers, filtering and quality control of cells used in scRNA sequencing analysis.** Related to Fig. 1 and Fig. 3.

**a**, table containing information on cell numbers, filter criteria for each batch of *in vitro* cultured embryos used for scRNA sequencing. **b**, UMI counts per cell across the different batches. **c**, Gene counts per cell across the different batches.

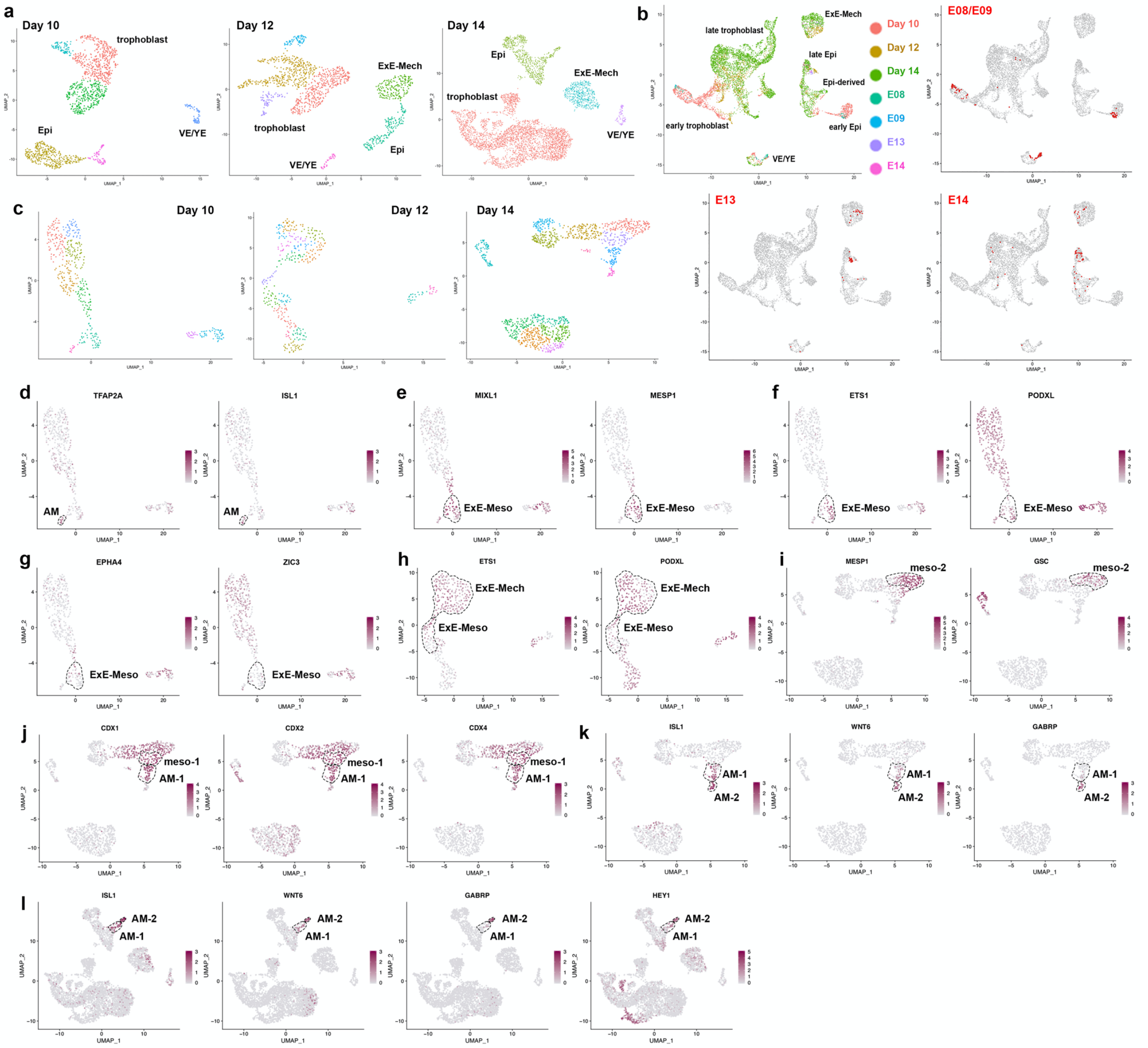


**Supplementary Fig. 3: Clustering, cell type identification and expression of marker genes from wildtype embryos.** Related to Fig. 1 and Fig. 2.

**a**, unsupervised clustering results of all cells from wildtype embryos used for cell type identification as shown in Fig. 1b. **b,** UMAP plot of the integrated dataset of all cells from the *in vitro* cultured embryos (Day 10, Day 12 and Day 14) with cells from the *in vivo* embryos (E08, E09, E13 and E14) colored by dataset and day corresponding to Fig. 1g. **c**, unsupervised clustering results of cells mapped to the epiblast (including its derivatives), endoderm or extraembryonic mesenchyme from embryos used for cell type identification as shown in Fig. 2a. **d-g**, Expression of marker genes in Day 10 embryos depicted on UMAP plots as shown in Fig. 2a. (**d**) amnion markers, (**e**) mesoderm markers, (**f**) extraembryonic mesoderm markers and (**g**) embryonic mesoderm markers. **h**, expression of markers for extraembryonic mesoderm in Day 12 embryos depicted on UMAP plot as shown in Fig 2a. **i-l**, expression of markers genes in Day 14 embryos depicted on UMAP plot as shown in Fig. 2a and 1b. (**i-j**) mesoderm markers and (**k-l**) amnion markers.

Epi, epiblast; Endo, endoderm; ExE-Meso, extraembryonic mesoderm; ExE-Mech, extraembryonic mesenchyme; AM-1, amnion 1; AM-2, amnion 2; meso-1, mesoderm 1; meso-2, mesoderm 2.

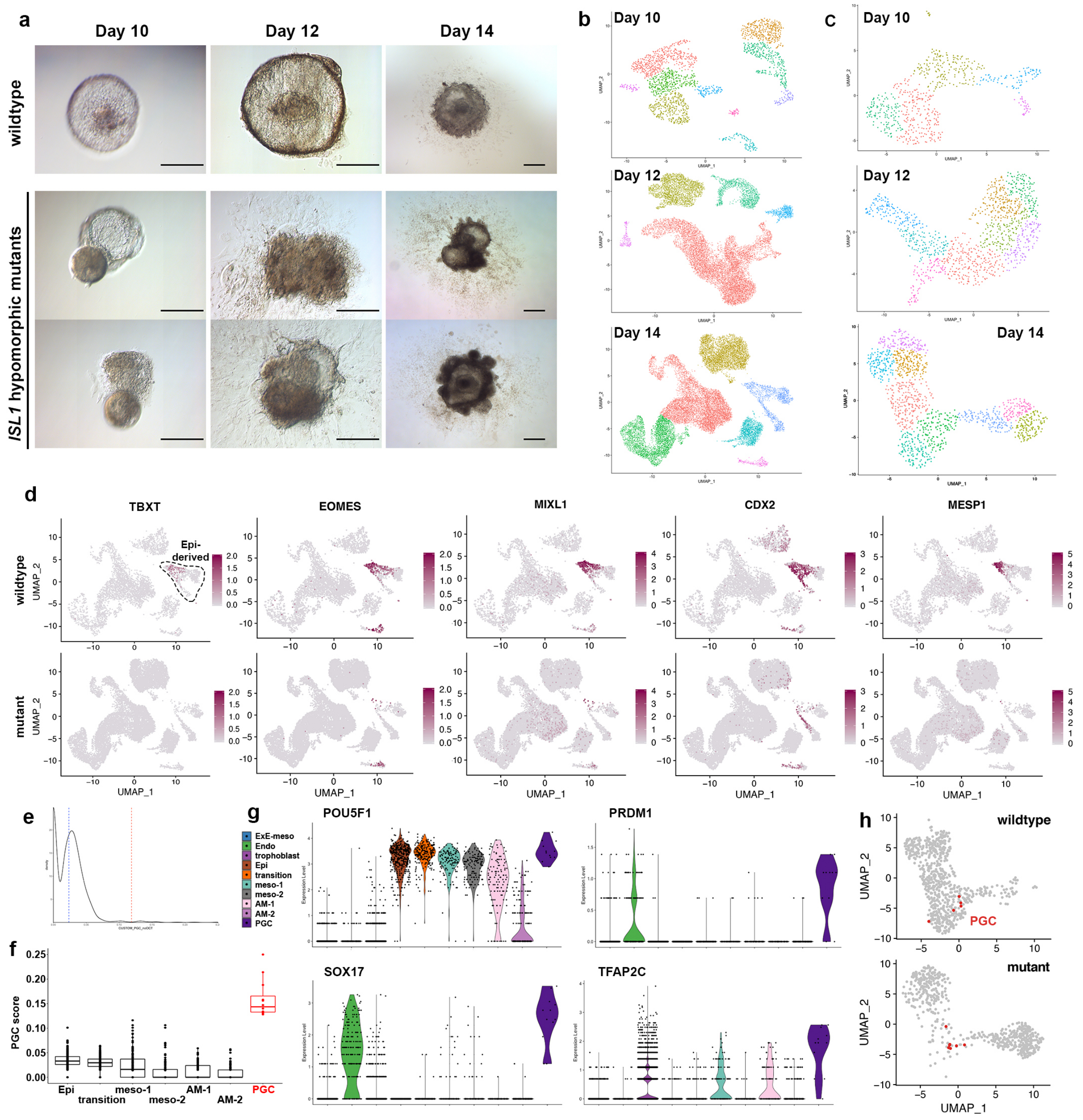


**Supplementary Fig. 4: Clustering of cells from the integrated analysis and gene expression in amnion of wildtype and *ISL1* hypomorphic mutants**. Related to Fig. 3.

**a**, morphology of in vitro cultured NHP embryos **b**, UMAP plot of unsupervised clustering results of all cells from the integrated analysis of wildtype and mutant embryos used for cell type as shown in Fig. 3b. **c**, unsupervised clustering results of cells mapping to the epiblast (including its derivatives) from the integrated analysis of wildtype and mutant embryos used for cell type identification as shown in Fig. 3c. **d**, expression of mesoderm markers in all cells from Day 14 embryos. **e**, density plot of the distribution of PGC scores across all cells from the epiblast and derivatives; blue line marks mean of distribution, red line marks 4 times standard deviation from mean. **f**, box plots of PGC scores in cells of the epiblast and its derivatives; PGCs marked in red. **g**, violin plot showing expression levels of selected PGC-specific genes across all cell populations in the dataset. **h**, UMAP plot of cells from the epiblast and its derivatives shown separately for cells from wildtype and mutant embryos; PGCs highlighted in red.

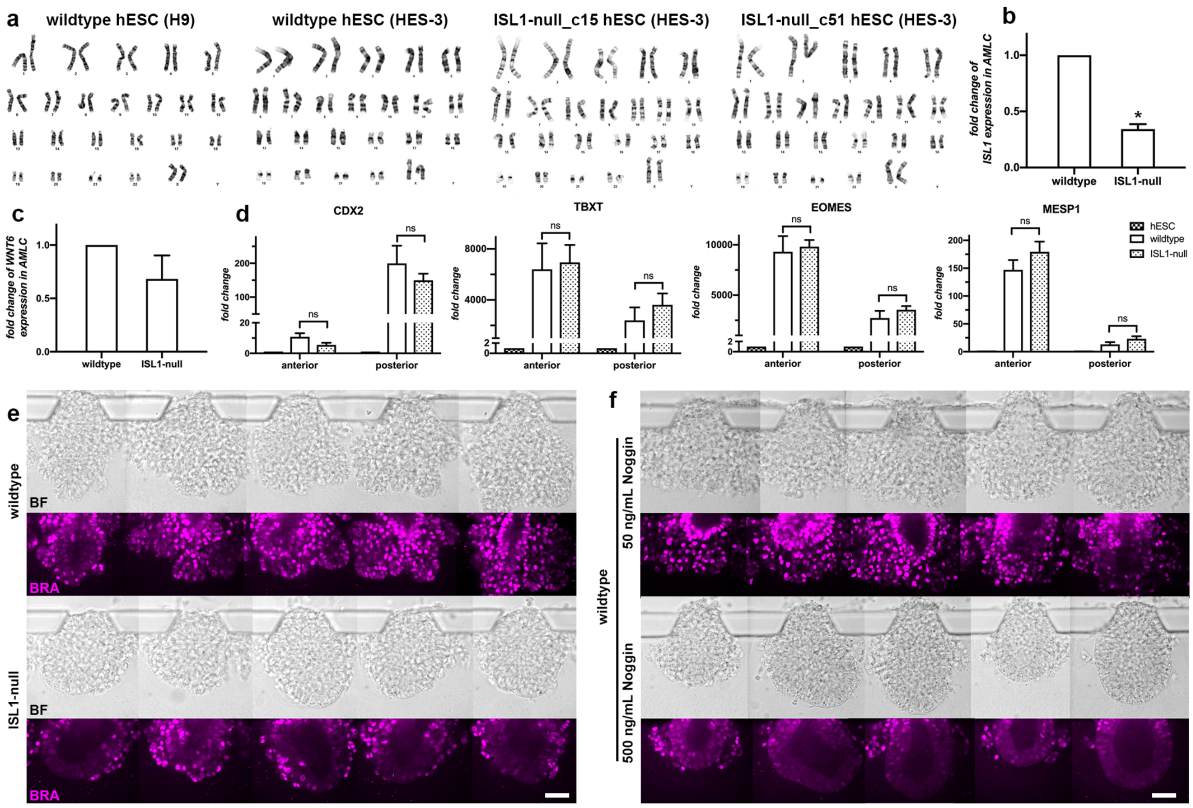


**Supplementary Fig. 5: *in vitro* assay using hESCs.** Related to Fig. 4.

**a**, karyotyping of the wildtype and the *ISL1*-null hESCs. **b**, expression of *ISL1* (p-value < 0.0001) in wildtype and ISL1-null AMLCs. **c**, expression of *BMP4* (p-value 0.0752) in wildtype and ISL1-null AMLCs. **d**, the expression of marker genes of anterior (*TBXT*, *EOMES*, *MESP1*) and posterior (*CDX2*) primitive streak at 40h after induction from wildtype and *ISL1*-null. Analyzed by student's t-test, all p-values > 0.05. Wildtype n=3; *ISL1*-null are aggregated values from two knockout cell lines n=3 for each. **e**, the morphology and the expression of BRA (magenta) in embryonic-like sacs derived from wildtype (upper) and *ISL1*-null (lower) at 48 hours after induction. **f**, the morphology and the expression of BRA (magenta) in embryonic-like sacs derived from wildtype treated with low (upper, 50 ng/mL) and high (lower, 500 ng/mL) dose of Noggin at 48 hours after induction. Scale bar 50 μm, n = 30 for each.
